# Germination response of invasive plants to soil burial depth and litter accumulation is species specific

**DOI:** 10.1101/2020.01.13.904227

**Authors:** Judit Sonkoly, Orsolya Valkó, Nóra Balogh, Laura Godó, András Kelemen, Réka Kiss, Tamás Miglécz, Edina Tóth, Katalin Tóth, Béla Tóthmérész, Péter Török

## Abstract

**Questions:** Plant invasions are considered among the biggest threats to biodiversity worldwide. In a full-factorial greenhouse experiment we analysed the effect of soil burial depth and litter cover on the germination of invasive plants. We hypothesised that (i) burial depth and litter cover affect the germination of the studied species, (ii) the effects of burial and litter cover interact with each other, and (iii) the effects are species-specific, but dependent on seed size.

**Methods:** We tested the germination and seedling growth of 11 herbaceous invasive species in a full-factorial experiment using four levels of seed burial depths and litter cover. We analysed the effect of burial, litter cover, and their interactions on germination, seedling length and biomass across species and at the species level.

**Results:** Soil burial depth and litter cover had a significant effect on the germination of the studied species, but there were considerable differences between species. We observed a general trend of species with bigger seeds being not or less seriously affected by soil burial and litter cover than smaller-seeded species. Correlations between seed weight and effect sizes mostly confirmed this general trend, but not in the case of soil burial.

**Conclusions:** Our findings confirmed that seed size is a major driver of species’ response to litter cover and to the combined effects of litter cover and soil burial, but there is no general trend regarding the response to soil burial depth. Despite its very small seeds the germination of *Cynodon dactylon* was not affected by soil burial. The germination of *Ambrosia artemisiifolia* was hampered by both soil burial and litter cover despite its relatively large seeds. Thus, specific information on species’ response to burial depth and litter accumulation is crucial when planning management or restoration in areas threatened by plant invasions.

## Introduction

The spread of invasive species is considered among the biggest threats to biodiversity worldwide (Early et al., 2016; Mollot, Pantel, & Romanuk, 2017). Invasive plants regularly colonize habitats like old-fields (Cramer, Hobbs, & Standish, 2008; Kelemen et al., 2016), whose area is increasing in several parts of the world due to the cessation of agricultural use and management (Prach, Török, & Bakker, 2017; Ramankutty & Foley, 1999). Perennial invasives can also persist until the later stages of succession, so they can even disrupt natural vegetation assembly and the recovery of old-fields (Cramer et al., 2008). The pathway of succession and species assembly can be ameliorated by restoration, but the presence of invasive plants can seriously hamper these projects as well (Cramer et al., 2008). Thus, where invasive species are a problem for the successful restoration of native communities, the most effective way can be the eradication of the viable propagules of invasive species from the sites, for example by limiting their germination and establishment using adequate management regimes (Regan, McCarthy, Baxter, Dane Panetta & Possingham, 2006).

Seed germination requires specific environmental conditions, so that the developing seedling has a higher chance of survival and establishment (Vandelook, Van der Moer, & Van Assche, 2008). Environmental factors like temperature, light and water availability, and the chemical environment affect seed germination (Baskin & Baskin, 1998), all of which substantially vary with soil depth (Benvenuti, Macchia, & Miele, 2001; Burmeier, Donath, Otte, & Eckstein, 2010). The depth from which a seedling can emerge depends on the amount of its seed reserves, which is mostly determined by the size of the seed (Humphries, Cauhan, & Florentine, 2018). Thus, the depth from which a seedling can emerge strongly depends on the size of the seed (Grundy, Mead, & Burston, 2003; Nandula, Eubank, Poston, Koger, & Reddy, 2006). Smaller seeds need to precisely track whether they are on or near the soil surface, for which their germination is most usually light-dependent (Burmeier et al., 2010; Milberg, Andersson, & Thompson, 2000). Seedlings of large-seeded species can emerge from greater depths, so their germination is typically not light-dependent (Jankowska-Blaszczuk & Daws, 2007; Milberg et al., 2000). Experimental studies demonstrated that germination and/or emergence rates generally decline with seed burial depth (e.g., Benvenuti et al., 2001; Burmeier et al., 2010; Traba, Azcaráte, & Peco, 2004), and that this relationship is often seed size-dependent (e.g., Bond, Honig, & Maze, 1999; Grundy et al., 2003; Limón & Peco, 2016). In turn, seed size and shape can strongly influence how easily seeds can be incorporated into deeper soil layers (Bekker et al., 1998), and also predict persistence in the soil (Thompson, Band, & Hodgson, 1993).

Besides soil burial depth, litter cover (i.e., the cover of dead plant material) is also a factor that can strongly affect seed germination and seedling establishment (Facelli & Pickett, 1991). The effects of litter on seedling establishment are mainly due to (i) the formation of a mechanical barrier (Donath & Eckstein, 2010), (ii) ameliorated water conditions (Eckstein, & Donath, 2005), (iii) decreased temperature fluctuations (Facelli & Pickett, 1991), (iv) decreased solar irradiation (Jensen & Gutekunst, 2003), (v) the changed red/far-red ratio of light (Jankowska-Blaszczuk & Daws, 2007), (vi) altered chemical environment by leached nutrients and allelochemicals (Ruprecht, Józsa, Ölvedi, & Simon, 2010), and (vii) protection from seed predators (Donath & Eckstein, 2012; Hölzel, 2005). Moreover, direct litter effects can also influence the outcome of plant-plant interactions, thus further altering the structure and composition of plant communities (Facelli & Pickett, 1991). Although the effects of litter on germination and establishment are mostly considered to be inhibitory (Jensen & Gutekunst, 2003; Eckstein & Donath, 2005), it can also have neutral and even facilitative effects depending on the amount of litter, the environmental conditions, and the species in question (Loydi, Eckstein, Otte, & Donath, 2013; Xiong & Nilsson, 1999). Several experimental results have confirmed the assumption that litter can have neutral or positive effects on seedling establishment, especially in the case of larger seeds (e.g. Hölzel, 2005; Myster, 2006; Miglécz, Tóthmérész, Valkó, Kelemen, & Török, 2013), when litter is present only in a small amount (Loydi et al., 2013; Xiong & Nilsson, 1999), or when water availability is suboptimal (Eckstein & Donath, 2005; Ruprecht et al., 2010).

Seedling establishment can be limited by the availability of seeds, the availability of suitable microsites, or both (Moore & Elmendorf, 2006). Therefore, if a site is already infested with the seeds of invasive species, the limitation of suitable microsites and ultimately the depletion of their viable seed bank can be an effective control method (Regan et al., 2006). Thus, to assess the risks posed by invasives and to plan management and restoration accordingly, information on their germination requirements is essential. There have been several studies dealing with their separate effects, but there are hardly any studies that simultaneously manipulated burial depth and litter cover. Rotundo & Aguiar (2005) found that litter had a positive effect on seedling establishment and growth when seeds were sown on the soil surface, but not when the seeds were buried to half the seed length. This suggests that the effects of litter cover and soil burial interact with each other, but more detailed studies are needed to unravel these effects. To fill this knowledge gap, our aim was to assess both the separate and the combined effects of soil burial and litter cover on the germination and establishment of invasive species. We set up a greenhouse experiment where 11 invasive species were sown with different soil burial depths and litter covers in a full-factorial design. We hypothesised that (i) soil burial and litter cover negatively affect the germination and establishment of the studied species, (ii) the effects of soil burial and litter cover interact with each other and that (iii) the effects are species-specific, but dependent on seed size.

## Materials & Methods

We selected 11 species for the experiment originating from different continents and acting as invasives in other regions of the world (Table 1). The species were selected to cover a high proportion of the seed size range of herbaceous invasive species. The studied species are mainly from the Asteraceae and Poaceae families, which corresponds to the fact that species from these families are disproportionately represented among the invasive plants for example in China (Weber, Sun, & Li, 2008) and in the Mediterranean regions (Andreu & Vilá, 2010), and that the Asteraceae and Poaceae families have the highest number of species in the global naturalised alien flora (Pyŝek et al., 2017). In addition, we selected *Asclepias syriaca*, as it is also colonising large areas of Europe and is especially widespread in Hungary (Tokarska-Guzik & Pisarczyk, 2015), representing a major threat in old-fields and degraded sand grasslands (Kelemen et al. 2016). Seeds were collected from natural populations in the Great Hungarian Plain (East Hungary, Central Europe) during 2016. Note that all seeds were collected in the same region where some of them are native and others are invasive aliens. We tested whether results differ between locally native and locally invasive species, and we found that there was no significant difference between the results of locally native and locally invasive species (Whelch Two-Sample t-tests for differences in the effect sizes of litter cover, soil burial and the two combined, *p*=0.399, *p*=0.526 and *p*=0.738, respectively). Henceforth we refer to the studied species as invasive species.

**Table 1.** Characteristics of the studied 11 species. TSW – Thousand-seed weight (g). Native and invasive alien ranges are given according to CABI Invasive Species Compendium (CABI, 2019) and the Invasive Plant Atlas of the United States (www.invasiveplantatlas.org). TSW values are based on Török et al. (2013).

| Species | Family | TSW | Native to | Alien invasive in |
| --- | --- | --- | --- | --- |
| *Conyza canadensis* | Asteraceae | 0.060 | North America | Europe |
| *Cynodon dactylon* | Poaceae | 0.114 | Africa | America, Oceania, Asia |
| *Solidago canadensis* | Asteraceae | 0.230 | North America | Europe, Asia, Oceania |
| *Lactuca serriola* | Asteraceae | 0.482 | Europe, Asia, North Africa | North America |
| *Cirsium arvense* | Asteraceae | 0.945 | Europe, Asia | North America |
| *Centraurea solstitialis* | Asteraceae | 1.366 | Europe, Asia, North Africa | North America |
| *Bromus tectorum* | Poaceae | 3.696 | Europe, Asia | North America |
| *Bromus inermis* | Poaceae | 4.574 | Europe, Asia | North America |
| *Ambrosia artemisiifolia* | Asteraceae | 5.656 | Central and North America | Africa, Asia, Australia, Europe |
| *Asclepias syriaca* | Asclepiadaceae | 6.063 | North America | Europe, Asia |
| *Tragopogon dubius* | Asteraceae | 6.529 | Europe, West Asia | North America |

We set up a greenhouse experiment in which we tested the effects of increasing seed burial depths (0 cm, 0.5 cm, 1 cm and 2 cm soil) and increasing levels of litter cover (0 g/m^2^, 150 g/m^2^, 300 g/m^2^ and 600 g/m^2^) an their interaction in a full-factorial design (altogether 16 treatments, Fig. 1). We filled pots with steam-sterilised potting soil and placed 25 seeds of a species on each of them, then applied soil burial and litter cover according to the treatment. We used the same potting soil for burying the seeds; litter was collected in a seminatural grassland dominated by the narrow-leaf grasses *Festuca rupicola* and *Poa angustifolia*. All 16 treatments were applied on all 11 species with 5 replications, meaning altogether 880 pots and 22,000 seeds. The pots were placed randomly in a greenhouse, frequently rearranged and watered daily with tap water to provide optimal conditions for germination. Germination lasted 6 weeks from the beginning of April 2017. At the end of the germination tests, we counted and removed all seedlings, measured the length of the shoot on 10 randomly chosen seedlings per pot (altogether 5,484 measurements) and weighed the aboveground biomass in each pot with an accuracy of 0.0001 g. Only those seedlings were counted that emerged above the soil surface, thus germinated successfully. Fatal germination events have not been counted.

**Figure 1.**
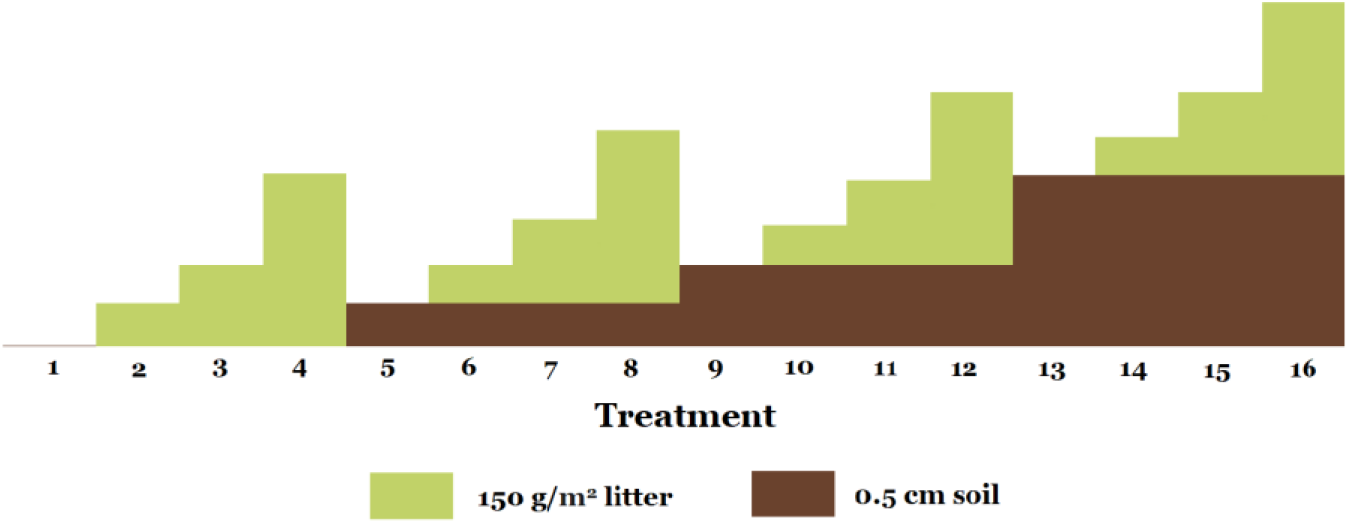
Soil burial depth and litter cover in the 16 treatments demonstrating the experimental design.

The effects of soil burial depth, litter cover, and their interactions on the standardized germination rate, standardized seedling length and standardized seedling biomass across species were analysed using two-way ANOVAs. The effects of soil burial depth, litter cover and their interactions on the species level were also analysed using two-way ANOVAs.

To demonstrate the separate effect of soil burial and litter cover, we compared standardized germination rates, seedling lengths and biomass in treatments 1, 2, 3 and 4, and treatments 1, 5, 9 and 13, respectively (see Fig. 1 for the treatments) using one-way ANOVA and Tukey HSD tests. Standardization was done in order to avoid the confounding effect of species identity, by considering values of germination rate, seedling length and seedling biomass of every species in the control treatment (i.e., treatment 1) as being 1, and expressing every other germination rate, seedling length and biomass value of the given species compared to the value obtained in the control. Thus, standardized values <1 represent a decrease, while values >1 represent an increase compared to the control.

To test if the effect sizes correlate with seed size, we calculated Cohen’s *d* (standardized mean difference, calculated as the difference of the means of control and treated groups divided by the weighted pooled standard deviations of these groups) for soil burial, litter cover and for the two combined for each species, and then used Spearman’s rank correlation tests. Thus, the standardized mean difference of treatment 1 (control) and treatment 4 was the effect size for litter cover; the standardized mean difference of treatment 1 and treatment 13 was the effect size for soil burial; and the standardized mean difference of treatment 1 and treatment 16 was the effect size for soil burial and litter cover combined (see Fig. 1).

*Conyza canadensis* and *Solidago canadensis* were excluded from the analyses of seedling length and biomass because of their very low (often 0%) germination rate in treatments other than the control. Some variables were log-transformed or square-root transformed to obtain normally distributed residuals. All statistical analyses were carried out in R statistical environment (R Development Core Team).

## Results

From the 22,000 seeds sown 9,480 seeds (43.1%) germinated. The mean germination rate of the studied species in the control treatment ranged from 0.11 (*Cynodon dactylon)* to 0.94 (*Bromus tectorum*), but the highest germination rate was not found in the control treatment in most species (except for *Centaurea solstitialis*, *Conyza canadensis* and *Solidago canadensis*) (Supplementary Figure 1).

Germination rate was significantly affected by both soil burial depth and litter cover, but not by their interactions, and both factors explained only a small fraction of the total variation (Table 2, Fig. 2).

**Table 2.** The effect of soil burial depth, litter cover and their interactions on the standardized germination rate across the studied species (two-way ANOVA). VC=variance component, i.e., the relative contribution of factors and their interactions to the total variation (expressed as the ratio of the sum of squares of the factor to the total sum of squares).

|  | df | *p* | VC |
| --- | --- | --- | --- |
| Soil burial depth | 3 | < 0.001 | 0.103 |
| Litter cover | 3 | < 0.001 | 0.035 |
| Soil × Litter | 9 | 0.833 | 0.005 |
| Residual | 864 |  | 0.857 |

**Figure 2.**
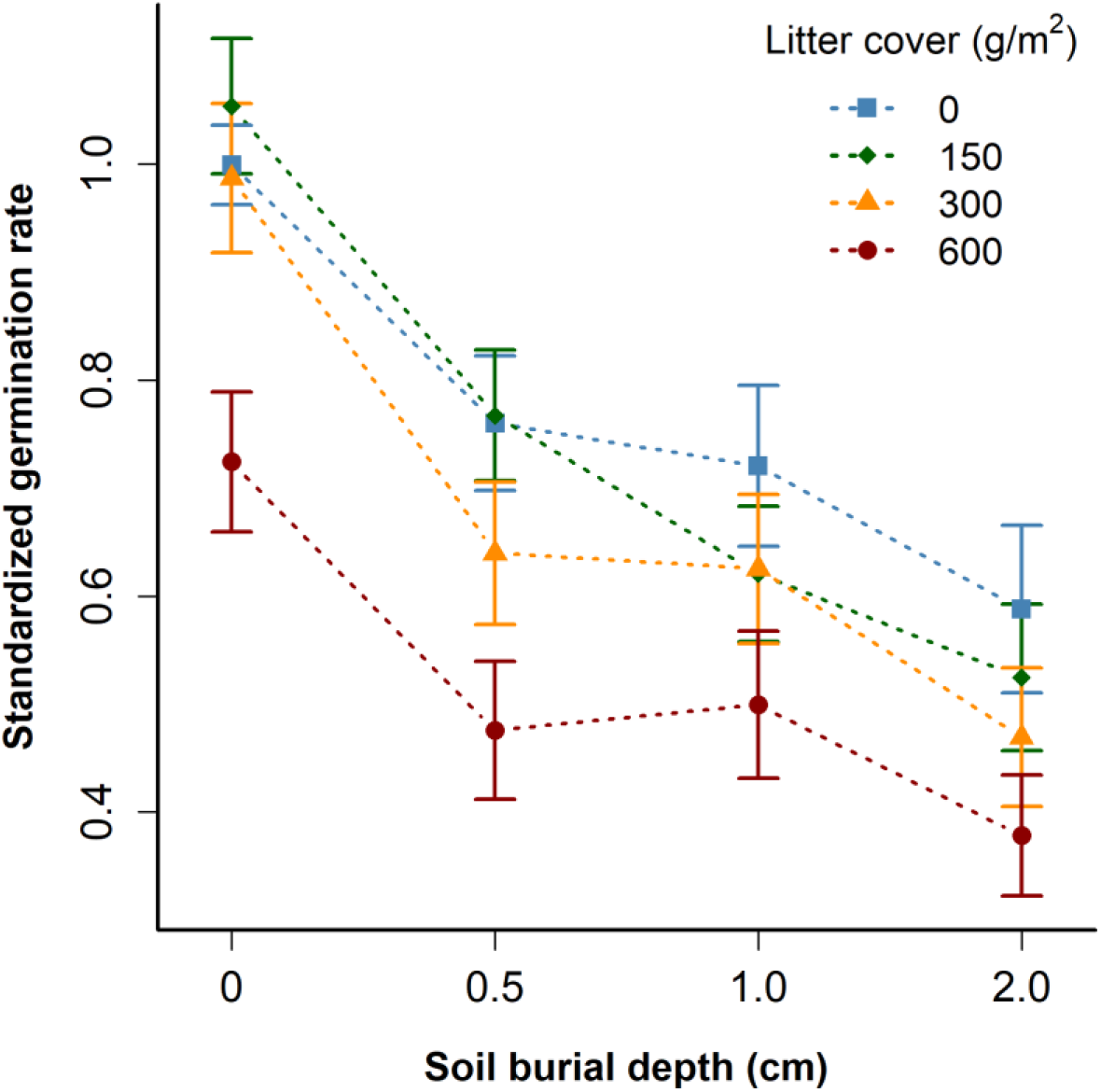
The effect of soil burial depth and litter cover on the standardized germination rate across the studied species (mean±SE).

Analysing the separate effects of soil burial and litter cover across species revealed that soil burial had a significant negative effect on the standardized germination rate (One-way ANOVA, F=7.019, *p*<0.001), and the difference was significant already at the lowest depth applied (0.5 cm, Fig. 3). Litter cover alone also had a significant negative effect on the standardized germination rate of species (One-way ANOVA, F=6.103, *p*<0.001), but the difference was significant only at the highest amount of litter applied (600 g/m^2^, Fig. 3).

**Figure 3.**
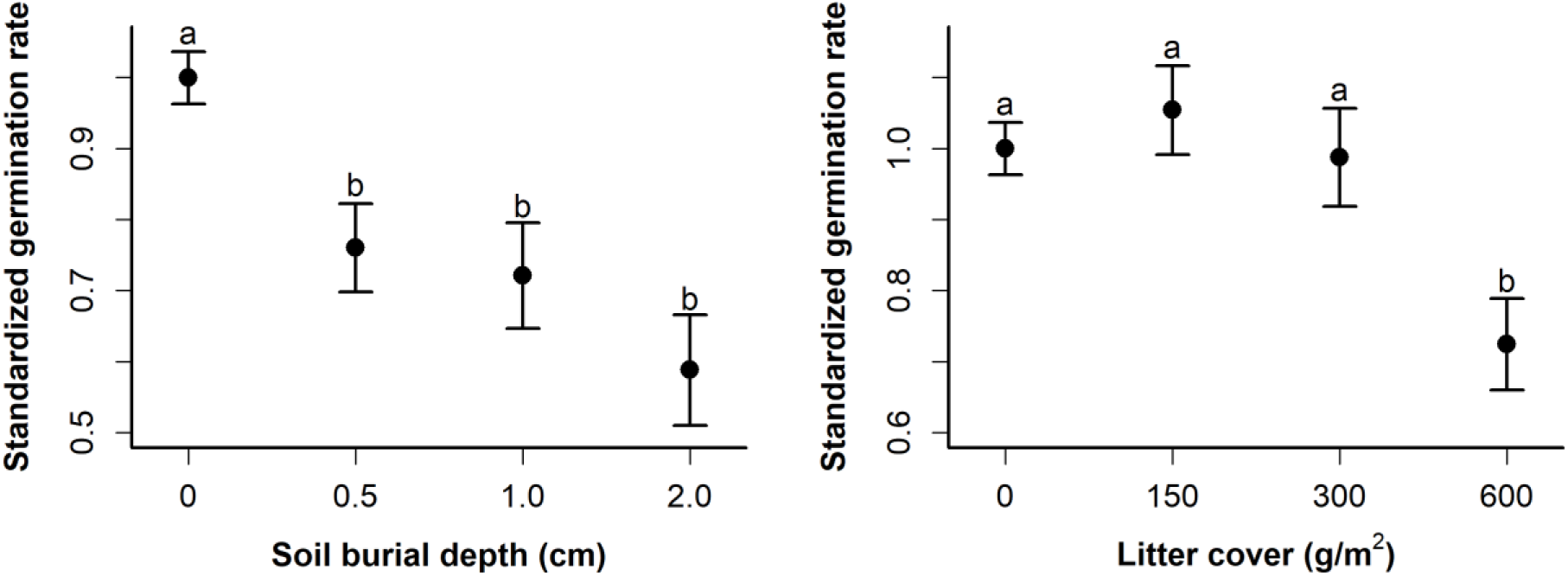
The separate effects of different levels of seed burial depth and litter cover across species. The standardized germination rates of species were compared between treatments 1, 5, 9 and 13 (for seed burial depth), and between treatments 1, 2, 3 and 4 (for litter cover), using one-way ANOVA and Tukey HSD tests. Different letters denote significant differences.

When analysed at the species level, the effects of both soil burial and litter cover were significant for most of the species, excluding some of the large-seeded ones (Table 3). The variance explained by soil burial was relatively high for the small-seeded species except for *Cynodon dactylon*. For the large-seeded species, the effect of seed burial depth was either not significant or explained only a small fraction of the total variation, except for *Ambrosia artemisiifolia*. The variance explained by litter cover was rather low for all the species (Table 3). The interaction term was significant for only a few species, and the variance explained by it was low for most species, except for *Solidago canadensis*. The unexplained variance was more than 50% for most of the large-seeded species and *Cynodon dactylon* (Table 3).

**Table 3.** The effect of soil burial depth, litter cover and their interaction on the germination rate of the studied species (2-way ANOVAs). VC =variance component, i.e., the relative contribution of factors and their interactions to the total variation (expressed as the ratio of the sum of squares of the factor to the total sum of squares).

| Species | Soil |  | Litter |  | Soil×Litter |  | Resid. |
| --- | --- | --- | --- | --- | --- | --- | --- |
|  | *p* | VC | *p* | VC | *p* | VC | VC |
| *Ambrosia artemisiifolia* | < 0.001 | 0.434 | < 0.001 | 0.165 | 0.065 | 0.085 | 0.316 |
| *Asclepias syriaca* | 0.584 | 0.027 | 0.573 | 0.028 | 0.925 | 0.052 | 0.893 |
| *Bromus inermis* | 0.391 | 0.039 | 0.610 | 0.023 | 0.373 | 0.126 | 0.812 |
| *Bromus tectorum* | 0.119 | 0.065 | 0.308 | 0.039 | 0.028 | 0.217 | 0.679 |
| *Centaurea solstitialis* | < 0.001 | 0.561 | < 0.001 | 0.222 | 0.134 | 0.040 | 0.177 |
| *Cirsium arvense* | < 0.001 | 0.445 | 0.012 | 0.077 | 0.391 | 0.063 | 0.415 |
| *Conyza canadensis* | < 0.001 | 0.617 | < 0.001 | 0.153 | < 0.001 | 0.182 | 0.048 |
| *Cynodon dactylon* | 0.006 | 0.112 | < 0.001 | 0.277 | 0.337 | 0.086 | 0.526 |
| *Lactuca serriola* | < 0.001 | 0.653 | 0.022 | 0.036 | 0.006 | 0.090 | 0.221 |
| *Solidago canadensis* | < 0.001 | 0.476 | < 0.001 | 0.144 | < 0.001 | 0.312 | 0.067 |
| *Tragopogon dubius* | 0.001 | 0.177 | 0.045 | 0.079 | 0.085 | 0.150 | 0.593 |

Standardized seedling length was significantly affected by both soil burial depth and litter cover, but not by their interaction, and both factors explained only a small fraction of the total variation (Supplementary Table 1, Supplementary Figure 2). When analysed at the species level, the effect of litter cover on seedling length was significantly positive for all species, and it explained a relatively high amount (>40%) of the variance for several species (Supplementary Table 2). Standardized seedling biomass was significantly affected by litter cover, but not by soil burial depth and their interaction (Supplementary Table 3, Supplementary Figure 3). When analysed at the species level, seedling biomass was significantly affected by soil burial depth, litter cover or their interactions in only some of the species, and the unexplained variance was relatively high for all the species (Supplementary Table 4). The separate effects of soil burial and litter cover were positive on both the standardized seedling length (F=10.64, *p*<0.001 for soil burial and F=32.75, *p*<0.001 for litter cover) and the standardized seedling biomass across species (F=5.275, *p*=0.002 for soil burial and F=5.145, *p*=0.02 for litter cover) (Supplementary Figure 4).

The correlation between seed weight and the effect size of seed burial on the germination rate of species was not significant (Fig. 4a), but seed weight was strongly correlated to both the effect size of litter cover (Fig. 4b) and to the effect size of the combined effect of soil burial and litter cover (Fig. 4c).

**Figure 4.**
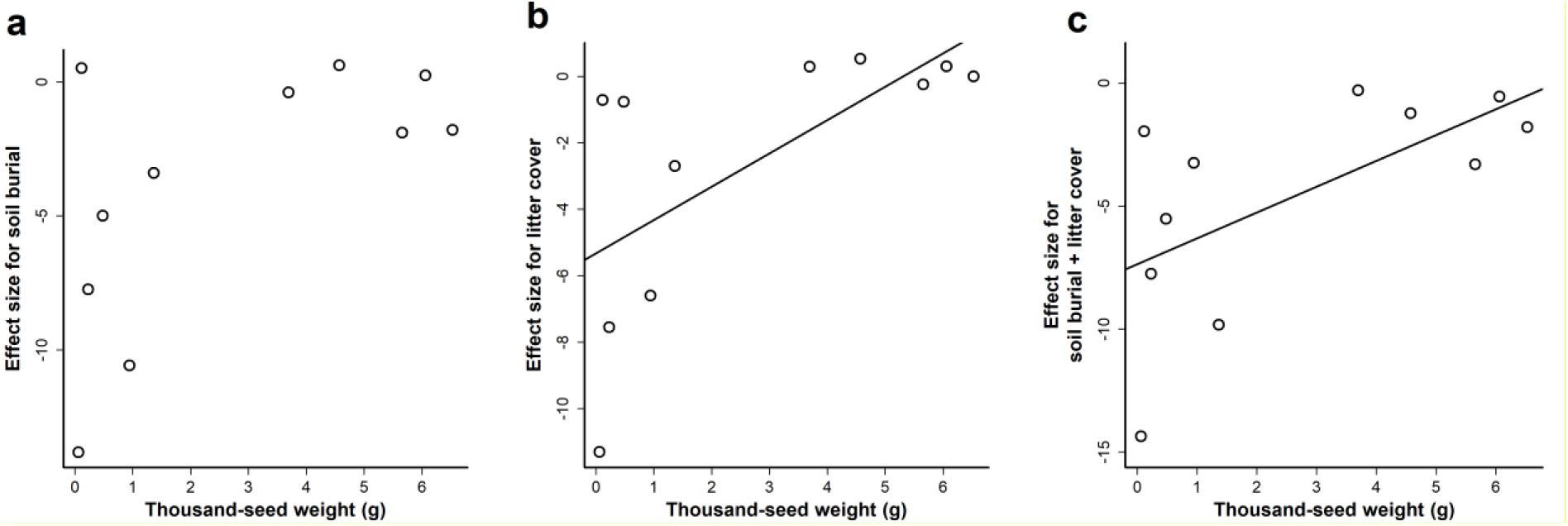
The correlation of seed weight and the effect size (Cohen’s *d*) for **(a)** soil burial (Spearman rank correlation, *p*=0.121), **(b)** litter cover (Spearman rank correlation, rho=0.764, *p*=0.009), and **(c)** litter cover and soil burial combined (Spearman rank correlation, rho=0.618, *p*=0.048) for the germination rate of species.

## Discussion

We hypothesised that soil burial and litter cover negatively affect the germination and establishment of the studied species. In our experiment, soil burial alone had a significant negative effect on the standardized germination rate of the studied species even at the lowest depth of soil burial applied (0.5 cm). This result agrees well with the findings of former experiments (e.g., Benvenuti et al., 2001; Burmeier et al., 2010; Traba et al., 2004). In turn, the standardized seedling length and biomass of the species were positively affected by soil burial; the seedlings grew bigger even at a 0.5 cm burial depth than with no burial, and the effect of burial was constant at every burial depth. This positive effect is probably due to the more optimal moisture conditions in deeper soil layers (Boyd & van Acker, 2003; Padilla & Pugnaire, 2007), which may have allowed seedlings that could germinate from deeper soil layers to grow taller and heavier than seedlings emerging directly from the soil surface.

Litter cover alone only had a significant negative effect on the standardized germination rate of the studied species when applied at the highest amount (600 g/m^2^). This finding is in accordance with former studies stating that litter cover can be advantageous for germination when it is present in a moderate amount or under certain conditions (Eckstein & Donath, 2005; Ruprecht et al., 2010). In a meta-analysis, Xiong & Nilsson (1999) demonstrated that small amounts of litter (< 200 g/m^2^) can even favour establishment and above-ground biomass. Similarly to soil burial, litter cover had a significant positive effect on standardized seedling length and standardized seedling biomass, probably because seedlings that were able to emerge below deeper litter layers were later able to grow taller and heavier due to the more favourable moisture conditions. It is also possible that litter has already started to decompose during the experiment and the seedlings benefited from the nutrient enrichment (Facelli & Pickett, 1991).

Previous studies have also analysed the separate effects of soil burial and litter cover on the germination of seeds including some invasive species as well (e.g., Guillemin & Chauvel, 2011; Reader, 1993), but there are hardly any results on their combined effects. Rotundo & Aguiar (2005) analysed the effects of litter on the seed germination of *Bromus pictus* in a greenhouse experiment, where seeds were positioned on the soil surface or buried in it to half the seed length. Their results indicated that litter increases the germination and emergence rates of surface-lying seeds, but not the emergence of buried seeds. The novelty of our study is that we simultaneously manipulated soil burial depth and litter cover, and we also tested whether the effects of soil burial and litter cover interact with each other. When analysed on the species level, the effect of the seed burial depth × litter cover interaction on the germination was significant for some of the species. This was probably due to deep soil burial masking the potential effects of litter cover, resulting in smaller or no effect of litter cover in treatments with deeper soil burial for the species *Cynodon dactylon*, *Lactuca serriola* and *Solidago canadensis* (see Fig. S1). The effect of the interaction term was significant for the length and the biomass of seedlings in even less of the studied species. These findings suggest that the effects of soil burial and litter cover interact with each other in some species, but this is not a general phenomenon.

We also hypothesised that the effects are species-specific, but dependent on seed size. When analysed across species, the effects of soil burial and litter cover on the standardized germination rate were significant, but their interaction was not, and the unexplained variance was high. This is well in accordance with the notion that the germination response to litter is dependent on species identity (Eckstein & Donath, 2005). At the species level, soil burial was significant for most of the species and it explained a relatively high amount of the variance in germination for several species. In the case of smaller-seeded species, variance explained by soil burial depth was relatively high, except for *Cynodon dactylon*, while in the case of larger-seeded species soil burial depth was either not significant or explained little variance, except for *Ambrosia artemisiifolia*, where soil burial explained a relatively high amount of the variance.

Our species-level results were mostly in line with former results about these species. Regarding *Asclepias syriaca*, Yenish, Fry, Durgan, & Wyse (1996) found that soil burial up to 2 cm did not affect its emergence, which is in accordance with our results. Our findings for the two *Bromus* species were quite similar; neither soil burial nor litter cover had a significant effect on their germination success. Prostko, Wu, Chandler, & Senseman (1997) observed no decline in the emergence of *B. tectorum* seeds from less than 5 cm soil burial depths, which corresponds to our results. Wicks, Burnside, & Fenster (1971) also stated that *B. tectorum* seeds germinated well under up to 2.5 cm (1 inch) of soil burial. In agreement with our findings, the field germination rate of *Centaurea solstitialis* was also found to be higher at a depth of 1 cm than at 2 cm (Larson & Kiemnec, 1997). We found that the germination of *Cirsium arvense* was strongly inhibited by soil burial, which was in accordance with the former findings that a burial of 1.5 cm (Tiley, 2010) and 2-3 cm (Laubhan & Shaffer, 2006) can inhibit its germination. By contrast, others indicate that *C. arvense* can emerge even from a depth of 6 cm (Wilson, 1979). Our results regarding *Conyza canadensis* were in accordance with Weaver (2001) and Nandula et al. (2006), who both reported that its emergence is maximum on the soil surface and decreases quickly with burial depth. In case of *Solidago canadensis*, we found that both soil burial and litter cover significantly reduce germination, and the inhibiting effect of litter cover for the germination of this species was also demonstrated by Goldberg & Werner (1983) and by Reader (1993). Our result that *Tragopogon dubius* was significantly negatively affected by soil burial depths up to 2 cm disagrees with Qi & Upadhyaya (1993), who found a high emergence rate of *T. dubius* at 2 cm burial depth.

We also analysed whether the separate and combined effects of soil burial and litter cover depend on seed size by testing the correlation of the effect size and the seed weight of each species. The effect sizes for the combined effects and for the separate effect of litter were strongly correlated with seed weight. Species with larger seeds had effect sizes close to zero, while the effect sizes for the small-seeded species were more variable, but in general larger (in the negative range). Our finding that the germination of smaller seeds is hampered by a litter layer of 600 g/m^2^ corresponds to the notion that smaller seeds absorb the amount of water needed for germination more rapidly than larger seeds (Kikuzawa & Koyama, 1999), so the positive effect of litter on moisture conditions is more pronounced in case of larger seeds, while small seeds, whose germination is usually light-dependent, are hampered by a thick litter layer (Milberg et al., 2000). In case of the separate effects of soil burial, the correlation of effect size with seed weight was not significant. Large-seeded species had effect sizes close to zero in this case as well, but for small-seeded species the effect sizes had an even broader range. From the small-seeded species, the response of *Cynodon dactylon* markedly differed from the others, having a positive effect size for soil burial. Having species whose germination behaviour deviate from the general trend corroborates the assumption that the light-dependence of germination is not simply determined by seed size, as it can also be species-specific and altered by adaptations to specific habitat conditions (Donath & Eckstein, 2005; Milberg et al., 2000).

There were two species that considerably deviated from the general trends that we identified. For other large-seeded species the effect of soil burial depth was either not significant or explained only a small amount of the variance in germination, but the germination of *Ambrosia artemisiifolia* was significantly affected by soil burial, and it also explained a relatively high proportion of the variation (see also Fig. S1). Former studies have also found that the germination of *A. artemisiifolia* seeds is decreasing with burial depth, being the highest on the soil surface (Essl et al., 2015; Guillemin & Chauvel, 2011). On this basis, soil tillage after high seed production is a feasible control option for *A. artemisiifolia* if it is not followed by deep soil tillage in the following year (Guillemin & Chauvel, 2011). Our results indicated that litter cover may also decrease the establishment of this species, but it is much less effective compared to soil burial. The germination of *A. artemisiifolia* being hampered by both soil burial and litter cover besides its relatively large seeds is most probably an adaptation to establishment on disturbed sites (Milberg et al., 2000).

The other species that refuted the expectations was *Cynodon dactylon*, as its germination was basically not affected by soil burial depth (see Appendix S1 and Table 3), despite having the second smallest seeds among the studied species. We suggest that the reason behind this germination behaviour may be that *C. dactylon* is a highly drought-tolerant species that regularly grows in very dry habitats (Shi, Wang, Cheng, Ye, & Chan, 2012), but its germination and/or seedlings may not be drought-tolerant. Thus, its seeds need to be able to germinate from greater depths where soil moisture conditions are more favourable. This assumption is in accordance with the finding of Mahmood, Malik, Lodhi, & Sheikh (1996) that an increase in soil moisture content significantly increases the germination of *C. dactylon* seeds.

Another important notion is that the type of litter may also influence the results, as different types of litter may affect germination and establishment differently (Myster, 2006), but previous results suggest that these differential effects are probably due to physical rather than chemical processes (Peterson & Facelli, 1992). In our experiment the physical and chemical effects of litter could not be differentiated, but as these effects are not separated under real life field conditions either, we aimed to analyse their net effects. Apart from seed size, seed shape is another seed trait which may influence the germination response to soil and litter cover (Facelli & Pickett, 1991), but the relationship seems to be complex, and it is possibly species-specific (Grundy et al., 2003). However, in case of the studied species the effect of seed shape could not be meaningfully tested, because seed size and shape (seed shape index, Thompson et al., 2003) were not independent in the studied species (bigger seeds were in general more elongated, except for *A. artemisiifolia*). Results indicate that seedling geometry may also influence whether a seedling can penetrate the litter layer (Facelli & Pickett, 1991). Soil burial and litter cover may also affect the temporal dynamics of germination, ultimately affecting vegetation dynamics and succession (Facelli & Pickett, 1991; Facelli & Facelli, 1993); thus, how the timing of germination can be altered is probably another important aspect to test.

Understanding the germination behaviour and control options of invasive species is even more crucial in the light of climate change and land use changes, which is expected to facilitate their further spread and the rise of new invasive species as well (e.g., Cunze, Leiblein, & Tackenberg, 2013; Hellmann, Byers, Bierwagen, & Dukes, 2008). Information on the cropping system used before abandonment may also be essential (e.g., Cardina, Herms, & Doohan, 2002; Guglielmini & Satorre, 2004). The use of reduced tillage cropping systems is rapidly spreading, and in such cases weed seeds accumulate on the soil surface or in the uppermost layers, enabling rapid germination of most species (Mohler, 1993). In order to identify if (and what kind of) tillage could be recommended for the control of invasives after abandonment, it is essential to know the soil depth from which seedlings of the locally problematic species can emerge (Humphries et al., 2018). Litter can also have an important role in the control of invasives, as it has already been demonstrated that it can hamper the establishment of weeds in old-fields (Deák et al., 2011) and decrease the emergence of invasives such as *Solidago canadensis* (Goldberg & Werner, 1983) or *Flaveria bidentis* (Li et al., 2016). Thus, a deeper insight into the effects of litter on the germination of invasives can be valuable in practical nature conservation. Another practical consideration is the effect of hay transfer as a restoration measure (Eckstein & Donath, 2005), as different amounts and types of litter can have differential effects on target species as well. Whether these effects differ between invasive and non-invasive species cannot be assessed based on our study. Contrary to our results on invasive plants, the germination response of non-invasive species to soil burial was found to be correlated with seed size in previous studies (Bond et al., 1999; Limón & Peco, 2016). The combined effects of soil burial and litter cover have only been studied on a non-invasive species previously, in which case their effects strongly interacted with each other (Rotundo & Aguiar, 2005).

Based on our results, the germination of most invasive plants is strongly affected by soil burial and litter cover, which limits their ability to establish on undisturbed natural or semi-natural communities. We can conclude that seed size is a major driver of the germination response to litter cover and to the combined effect of litter and soil burial, but there is no general trend regarding the response to soil burial depth. From the studied species the germination of the small-seeded *Cynodon dactylon* was practically not affected by soil burial, while the much bigger seeds of *Ambrosia artemisiifolia* were significantly inhibited, which may be attributed to the influence of their habitat-preferences and strategy. Thus, to effectively deal with the ever-increasing threat of invasive plants, species-specific information on their germination response to litter cover and soil burial is essential. Such species-specific information would not only aid restoration but could also improve our ability to assess their risks posed on natural and semi-natural habitats. Furthermore, a more general implication is that although seed size is an important driver of plants’ response to several factors and it is associated with many plant attributes, other effects such as that of habitat preference or plant strategy can mask the influence of seed size.

## Author contribution

B.T., O.V. and P.T. designed the study; J.S., N.B., L.G., A.K., R.K., T.M., E.T., K.T. and O.V. carried out the experiments and collected the data; J.S. analysed the data; J.S. led the writing of the manuscript and all authors contributed critically to the drafts and gave final approval for publication.

## Acknowledgements

We would like to thank K. Lukács and S. Radócz for their help during the experiments.

## Notes

**Funding information** This project was funded by OTKA K 116639 (BT), NKFI KH 126477 (BT), NKFIH K 119225 (PT), NKFI FK 124404 (OV), NKFI KH 126476 (OV), NKFIH PD 124548 (TM), NKFIH KH 130320 (ET) and NKFI PD 128302 (KT). OV and AK were supported by the Bolyai János Research Scholarship of the Hungarian Academy of Sciences. AK was supported by MTA’s Postdoctoral Research Program.

