## Supplementary material for "Germination response of invasive plants to soil burial depth and litter accumulation is species specific"

**Supplementary Figure 1.** The germination rate of species in the applied treatments. Species are presented in the order of increasing seed weight. Treatments (litter covers and seed burial depths) are demonstrated in the lowest panels.

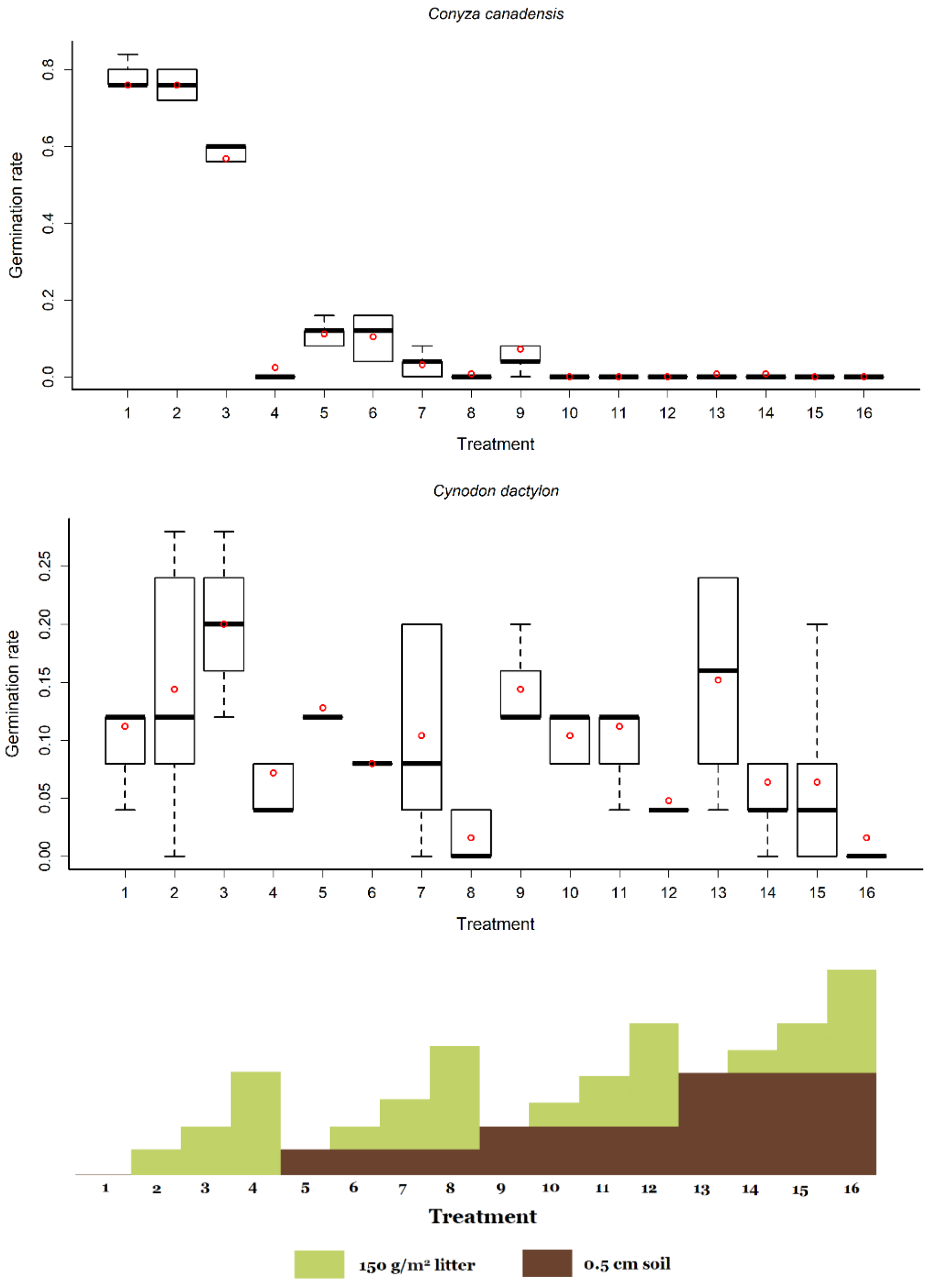

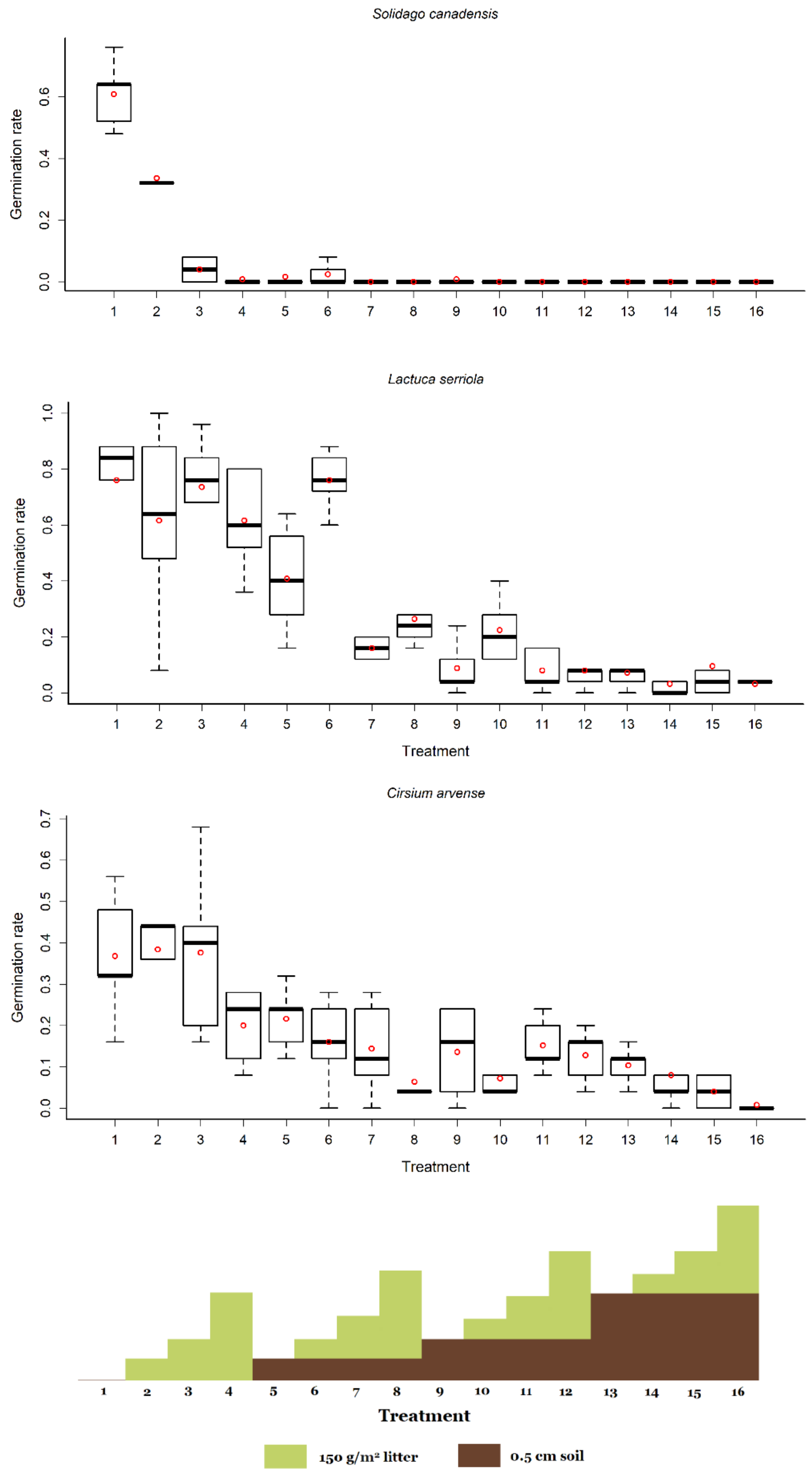
Solidago canadensis
Germination rate
0.0
0.2
0.4
0.6
1 2 3 4 5 6 7 8 9 10 11 12 13 14 15 16
Lactuca serriola
Germination rate
0.0
0.2
0.4
0.6
0.8
1.0
1 2 3 4 5 6 7 8 9 10 11 12 13 14 15 16
Treatment
Cirsium arvense
Germination rate
0.0
0.1
0.2
0.3
0.4
0.5
0.6
0.7
1 2 3 4 5 6 7 8 9 10 11 12 13 14 15 16
Treatment
1 2 3 4 5 6 7 8 9 10 11 12 13 14 15 16
Treatment
150 g/m² litter
0.5 cm soil

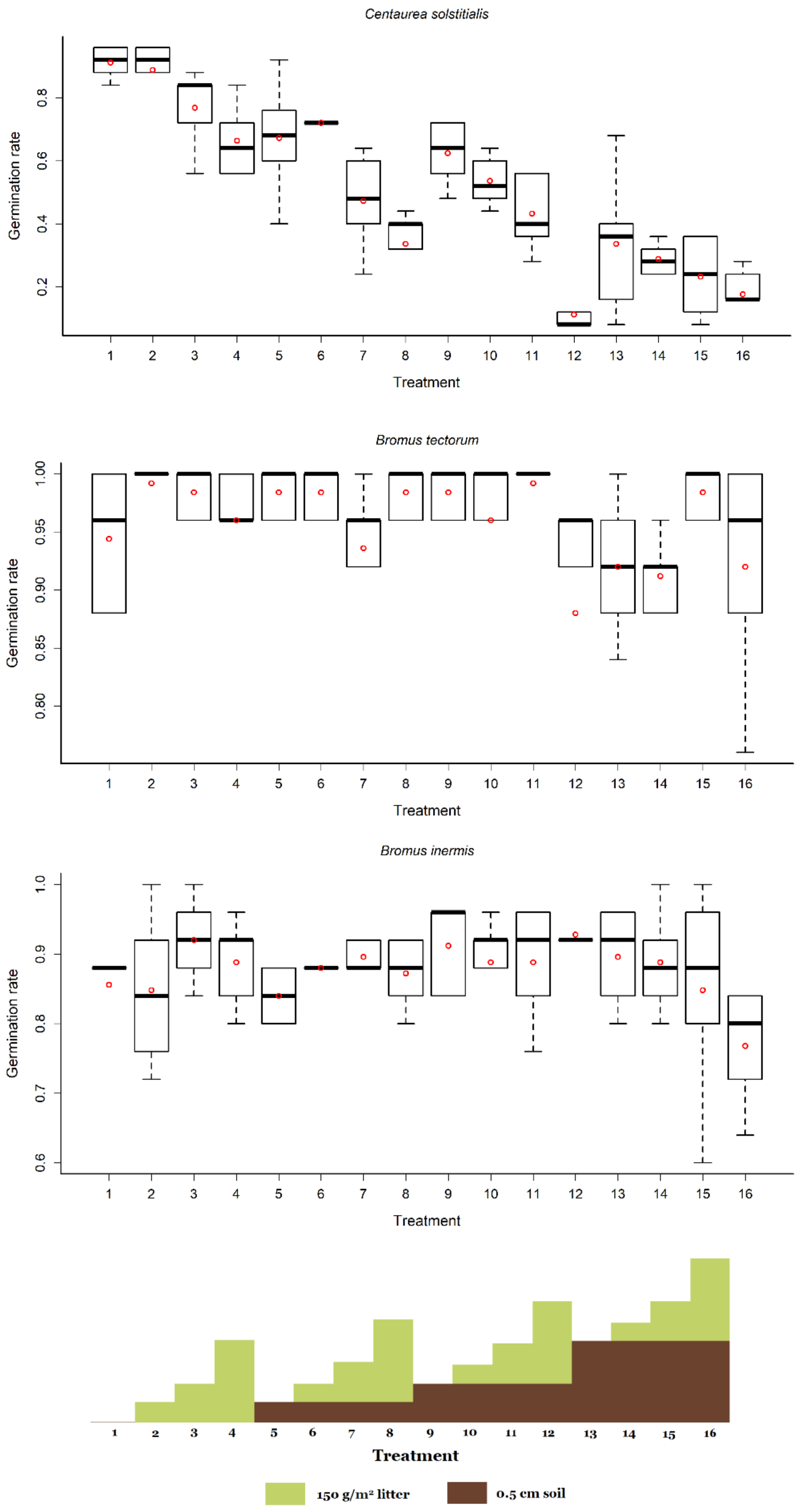

Centaurea solstitialis
Germination rate
0.2
0.4
0.6
0.8
1 2 3 4 5 6 7 8 9 10 11 12 13 14 15 16
Treatment
Bromus tectorum
Germination rate
0.80
0.85
0.90
0.95
1.00
1 2 3 4 5 6 7 8 9 10 11 12 13 14 15 16
Treatment
Bromus inermis
Germination rate
0.6
0.7
0.8
0.9
1.0
1 2 3 4 5 6 7 8 9 10 11 12 13 14 15 16
Treatment
1 2 3 4 5 6 7 8 9 10 11 12 13 14 15 16
Treatment
150 g/m² litter
0.5 cm soil

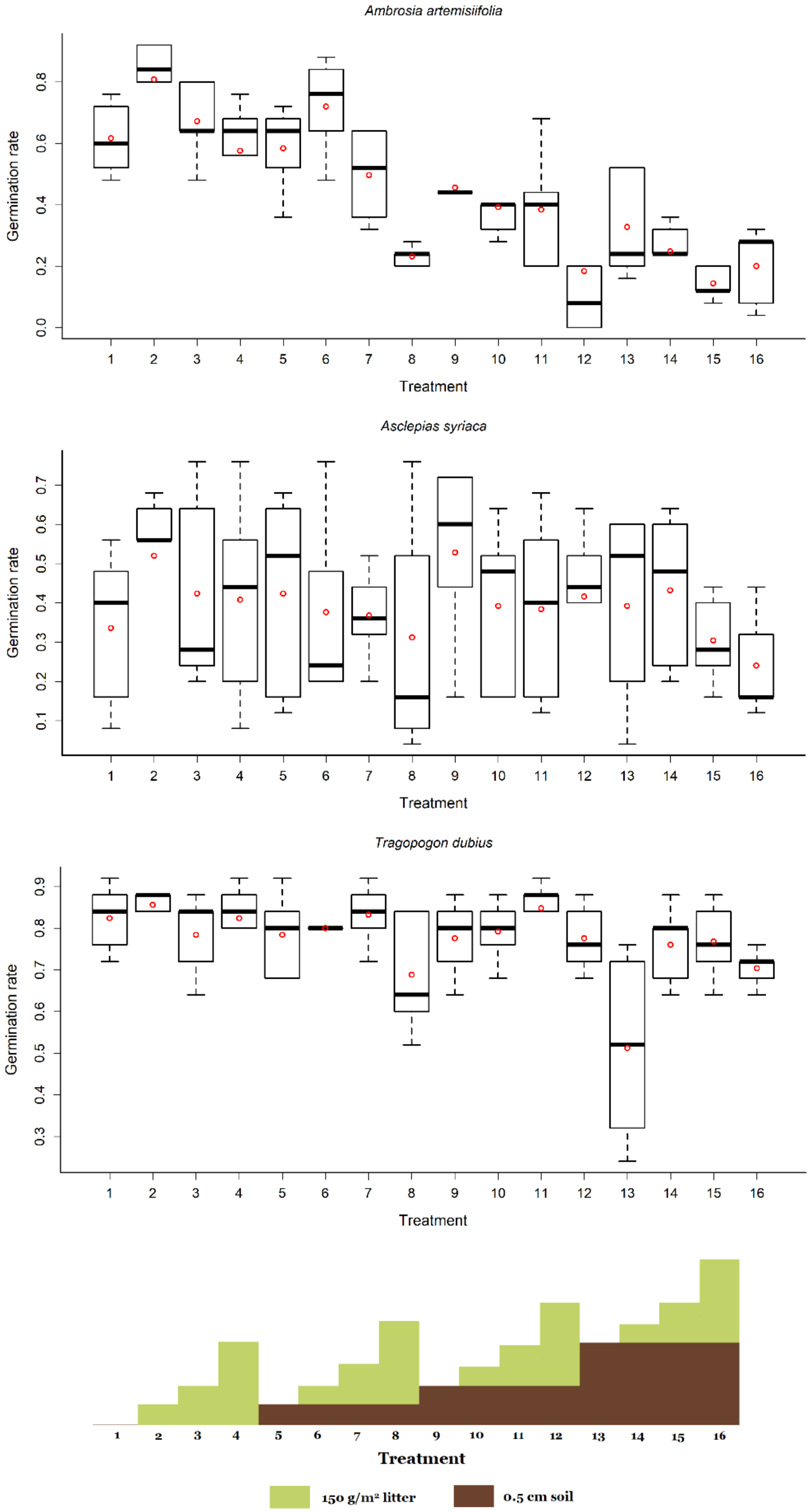
Ambrosia artemisiifolia
Germination rate
Treatment
Asclepias syriaca
Germination rate
Treatment
Tragopogon dubius
Germination rate
Treatment
Treatment
150 g/m² litter
0.5 cm soil

**Supplementary Table 1.** The effect of seed burial depth, litter cover, and their interactions on the standardized seedling length of across the studied species (two-way ANOVA). *Conyza canadensis* and *Solidago canadensis* were excluded from the analyses, because of their very low germination rates in treatments other than other than the control. Notations: VC =variance component, i.e., the relative contribution of factors and their interactions to the total variation (expressed as the ratio of the sum of squares of the factor to the total sum of squares).

| | df | *p* | VC |
|---|---|---|---|
| Soil burial depth | 3 | < 0.001 | 0.026 |
| Litter cover | 3 | < 0.001 | 0.128 |
| Soil × Litter | 9 | 0.070 | 0.020 |
| Residual | 667 | | 0.826 |

**Supplementary Figure 2.** The effect of soil burial depth and litter cover on the standardized seedling length across the studied species (mean±SE).

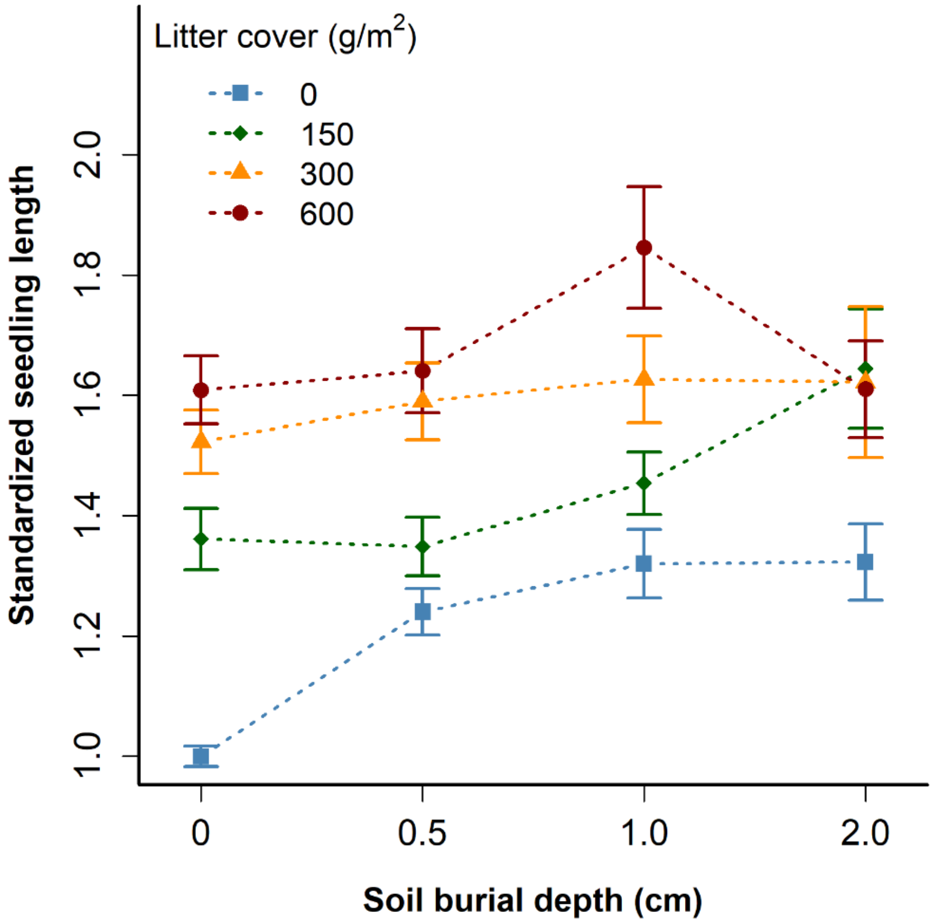

**Supplementary Table 2.** The effect of seed burial depth, litter cover, species identity and their interactions on the seedling length of the studied species (two-way ANOVAs). *Conyza canadensis* and *Solidago canadensis* were excluded from the analyses, because of their very low germination rates in treatments other than the control. VC =variance component, i.e., the relative contribution of factors and their interactions to the total variation (expressed as the ratio of the sum of squares of the factor to the total sum of squares).

| **Species** | **Soil** | | **Litter** | | **Soil×Litter** | | **Resid.** |
|---|---|---|---|---|---|---|---|
| | ***p*** | **VC** | ***p*** | **VC** | ***p*** | **VC** | **VC** |
| *Ambrosia artemisiifolia* | 0.114 | 0.070 | 0.006 | 0.155 | 0.676 | 0.075 | 0.701 |
| *Asclepias syriaca* | < 0.001 | 0.214 | 0.002 | 0.133 | 0.035 | 0.155 | 0.497 |
| *Bromus inermis* | < 0.001 | 0.233 | < 0.001 | 0.449 | 0.258 | 0.049 | 0.270 |
| *Bromus tectorum* | < 0.001 | 0.237 | < 0.001 | 0.437 | 0.001 | 0.108 | 0.217 |
| *Centaurea solstitialis* | 0.069 | 0.066 | < 0.001 | 0.289 | 0.502 | 0.075 | 0.570 |
| *Cirsium arvense* | 0.357 | 0.040 | 0.001 | 0.231 | 0.673 | 0.080 | 0.649 |
| *Cynodon dactylon* | 0.094 | 0.065 | < 0.001 | 0.343 | 0.360 | 0.098 | 0.493 |
| *Lactuca serriola* | 0.212 | 0.042 | 0.004 | 0.133 | < 0.001 | 0.353 | 0.472 |
| *Tragopogon dubius* | < 0.001 | 0.124 | < 0.001 | 0.454 | 0.467 | 0.051 | 0.371 |

**Supplementary Table 3.** The effect of seed burial depth, litter cover, and their interactions on the standardized seedling biomass of the studied species (2-way ANOVA). *Conyza canadensis* and *Solidago canadensis* were excluded from the analyses, because of their very low germination rate in treatments other than the control. VC =variance component, i.e., the relative contribution of factors and their interactions to the total variation (expressed as the ratio of the sum of squares of the factor to the total sum of squares).

| | **df** | ***p*** | **VC** |
|---|---|---|---|
| Soil burial depth | 3 | 0.282 | 0.006 |
| Litter cover | 3 | 0.005 | 0.019 |
| Soil × Litter | 9 | 0.130 | 0.020 |
| Residual | 665 | | 0.956 |

**Supplementary Figure 3.** The effect of soil burial depth and litter cover on the standardized seedling biomass across the studied species (mean±SE).

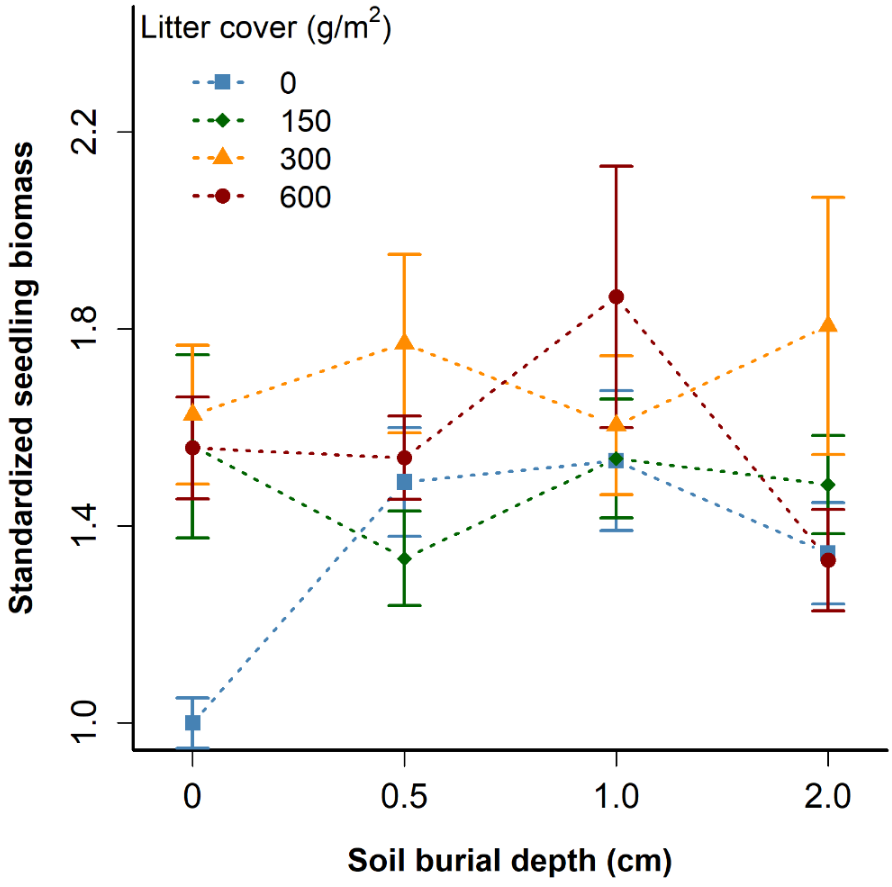


**Supplementary Table 4.** The effect of seed burial depth, litter cover and their interaction on the seedling biomass of the studied species (two-way ANOVAs). *Conyza canadensis* and *Solidago canadensis* were excluded from the analyses, because of their very low germination rates in treatments other than the control. VC =variance component, i.e., the relative contribution of factors and their interactions to the total variation (expressed as the ratio of the sum of squares of the factor to the total sum of squares).

| **Species** | **Soil** | | **Litter** | | **Soil×Litter** | | **Resid.** |
|---|---|---|---|---|---|---|---|
| | ***p*** | **VC** | ***p*** | **VC** | ***p*** | **VC** | **VC** |
| *Ambrosia artemisiifolia* | 0.443 | 0.038 | 0.813 | 0.013 | 0.654 | 0.094 | 0.855 |
| *Asclepias syriaca* | 0.005 | 0.140 | 0.110 | 0.063 | 0.078 | 0.166 | 0.631 |
| *Bromus inermis* | 0.152 | 0.055 | 0.001 | 0.190 | 0.264 | 0.115 | 0.641 |
| *Bromus tectorum* | 0.010 | 0.086 | < 0.0001 | 0.293 | 0.006 | 0.180 | 0.442 |
| *Centaurea solstitialis* | 0.003 | 0.132 | 0.001 | 0.155 | 0.024 | 0.179 | 0.534 |
| *Cirsium arvense* | 0.193 | 0.058 | 0.055 | 0.096 | 0.068 | 0.205 | 0.641 |
| *Cynodon dactylon* | 0.144 | 0.068 | 0.034 | 0.112 | 0.067 | 0.209 | 0.611 |
| *Lactuca serriola* | 0.979 | 0.003 | 0.956 | 0.005 | 0.044 | 0.267 | 0.726 |
| *Tragopogon dubius* | 0.060 | 0.068 | < 0.0001 | 0.280 | 0.310 | 0.094 | 0.558 |

**Supplementary Figure 4.** Separate effects of litter cover and seed burial depth across species. Standardized seedling lengths and standardized seedling biomass of species were compared between treatments 1, 2, 3 and 4 (for litter cover), and between treatments 1, 5, 9 and 13 (for seed burial depth), using one-way ANOVA and Tukey HSD tests. Different letters indicate significant differences between groups. Different letters denote significant differences.

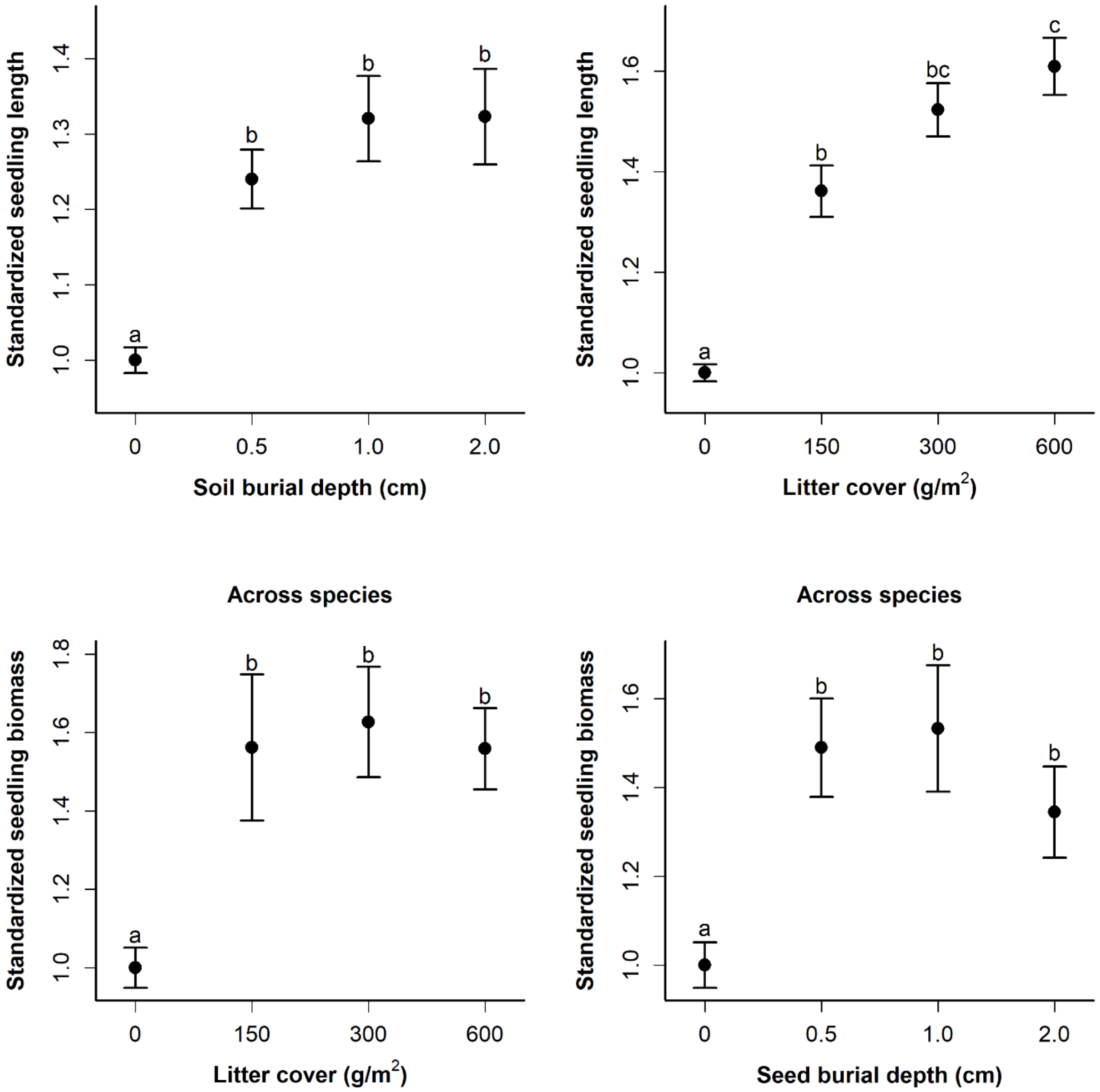
